# The Practical Haplotype Graph, a platform for storing and using pangenomes for imputation

**DOI:** 10.1101/2021.08.27.457652

**Authors:** PJ Bradbury, T Casstevens, SE Jensen, LC Johnson, ZR Miller, B Monier, MC Romay, B Song, ES Buckler

## Abstract

**Motivation:** Pangenomes provide novel insights for population and quantitative genetics, genomics, and breeding not available from studying a single reference genome. Instead, a species is better represented by a pangenome or collection of genomes. Unfortunately, managing and using pangenomes for genomically diverse species is computationally and practically challenging. We developed a trellis graph representation anchored to the reference genome that represents most pangenomes well and can be used to impute complete genomes from low density sequence or variant data.

**Results:** The Practical Haplotype Graph (PHG) is a pangenome pipeline, database (PostGRES & SQLite), data model (Java, Kotlin, or R), and Breeding API (BrAPI) web service. The PHG has already been able to accurately represent diversity in four major crops including maize, one of the most genomically diverse species, with up to 1000-fold data compression. Using simulated data, we show that, at even 0.1X coverage, with appropriate reads and sequence alignment, imputation results in extremely accurate haplotype reconstruction. The PHG is a platform and environment for the understanding and application of genomic diversity.

**Availability:** All resources listed here are freely available. The PHG Docker used to generate the simulation results is https://hub.docker.com/ as maizegenetics/phg:0.0.27. PHG source code is at https://bitbucket.org/bucklerlab/practicalhaplotypegraph/src/master/. The code used for the analysis of simulated data is at https://bitbucket.org/bucklerlab/phg-manuscript/src/master/. The PHG database of NAM parent haplotypes is in the CyVerse data store (https://de.cyverse.org/de/) and named /iplant/home/shared/panzea/panGenome/PHG_db_maize/phg_v5Assemblies_20200608.db.

## Introduction

Individuals within a species or population can vary considerably for genome content. A single reference genome often inadequately represents that diversity. As a result, a system for organizing and using information from multiple genomes would be very useful. Several groups have shown that pangenome graphs can effectively represent genomic diversity (Eizenga *et al*., 2020; Llamas *et al*., 2019; Rakocevic *et al*., 2019). Most of that work has been done with human genomes. Other species, especially plant species, have more complex and diverse genomes, with an order of magnitude more active transposons, making the direct application of pangenome graphs developed for humans difficult (Llamas *et al*., 2019). For example, comparisons of maize sequence estimate that 40% to 50% of the genome is not alignable between pairs of inbred lines (Brunner *et al*., 2005; Sun *et al*., 2018). In contrast, a study of 910 humans found that total unalignable sequence from all individuals equaled about 10% of the genome (Sherman *et al*., 2019) while an earlier study of an African and an Asian genome found about 0.2% of the genomic sequence of each individual absent from the human reference genome (Li *et al*., 2010).

The Practical Haplotype Graph (PHG) differs from other graphical approaches in the way that it handles larger scale variation. Specifically, because retroelements dominate many plant genomes (Baucom *et al*., 2009; Bennetzen, 2000), intergenic regions often align poorly between different individuals. In addition, plants generally have numerous but compact genes with recombination focused near them (Schnable *et al*., 1998; Rodgers-Melnick *et al*., 2015). The genes are broadly shared between individuals, while intergenic regions are far more divergent.

These features suggest dividing regions of the genome into genic and intergenic intervals. A second element, which arises from coalescent processes and from bottlenecks introduced by crop breeding, is that haplotypes for these intervals tend to be diverse but limited in number, especially within breeding programs.

This paper introduces the PHG, which represents genomes as sequences of haplotypes rather than nucleotides and uses a simplified approach for dealing with structural variation. The PHG encapsulates a relational database for storing sequence and alignment information, software for building and using that database, and pipelines that employ widely used third-party software for some of those tasks. The PHG software and database schemas are open-source and freely available. They are distributed as a Docker image to simplify the task of installing the environment needed.

While the PHG supports a range of genomic analyses, it is also designed to be a useful tool for breeding programs. As such it provides methods to impute genotypes based on the haplotypes stored in the database using low coverage sequence or single nucleotide polymorphisms (SNPs) as input. It also divides the genome into genic and intergenic haplotypes, which are biologically interesting units likely to have a limited number of common haplotypes.

The input sequence can be a whole genome sequence generated from a random shear library or sequence based on reduced representation libraries created by a variety of methods. Alternatively, SNP calls themselves can be used to impute the haplotypes carried by an individual, though most of the description below refers to the use of short read sequence to impute haplotypes and genotypes. That flexibility allows a user to choose a sequencing methodology based on cost and to change methods or suppliers over time as logistics dictate. It also provides a method for translating historical genotypes from various methods onto a common platform. Because the low coverage sequence is used to impute a path through the graph, it results in a list of imputed haplotypes. Storing the haplotype lists in the PHG database results in very compact storage of a large number of genotypes in a single database. Downstream analyses can either use VCF files of imputed nucleotide variants or use the haplotypes directly as multi-allelic loci.

## System and methods

The PHG is a trellis graph that represents discrete genomic DNA sequences and connections between them (Figure 1). A haplotype is defined as the sequence of part of an individual chromosome. A reference range is a segment of the reference genome. A node in the graph represents one haplotype for one reference range. Two physically adjacent nodes are connected by an edge, which does not have an associated sequence but only indicates whether two haplotypes (might) occur together in some individuals. If a graph is built from a subset of ranges, for example genic ranges only, it does not contain any information about the excluded ranges. A haplotype graph is then a collection of nodes and edges. Nodes for a single range can be thought of as multiple alleles at a single position.

**Figure 1.**
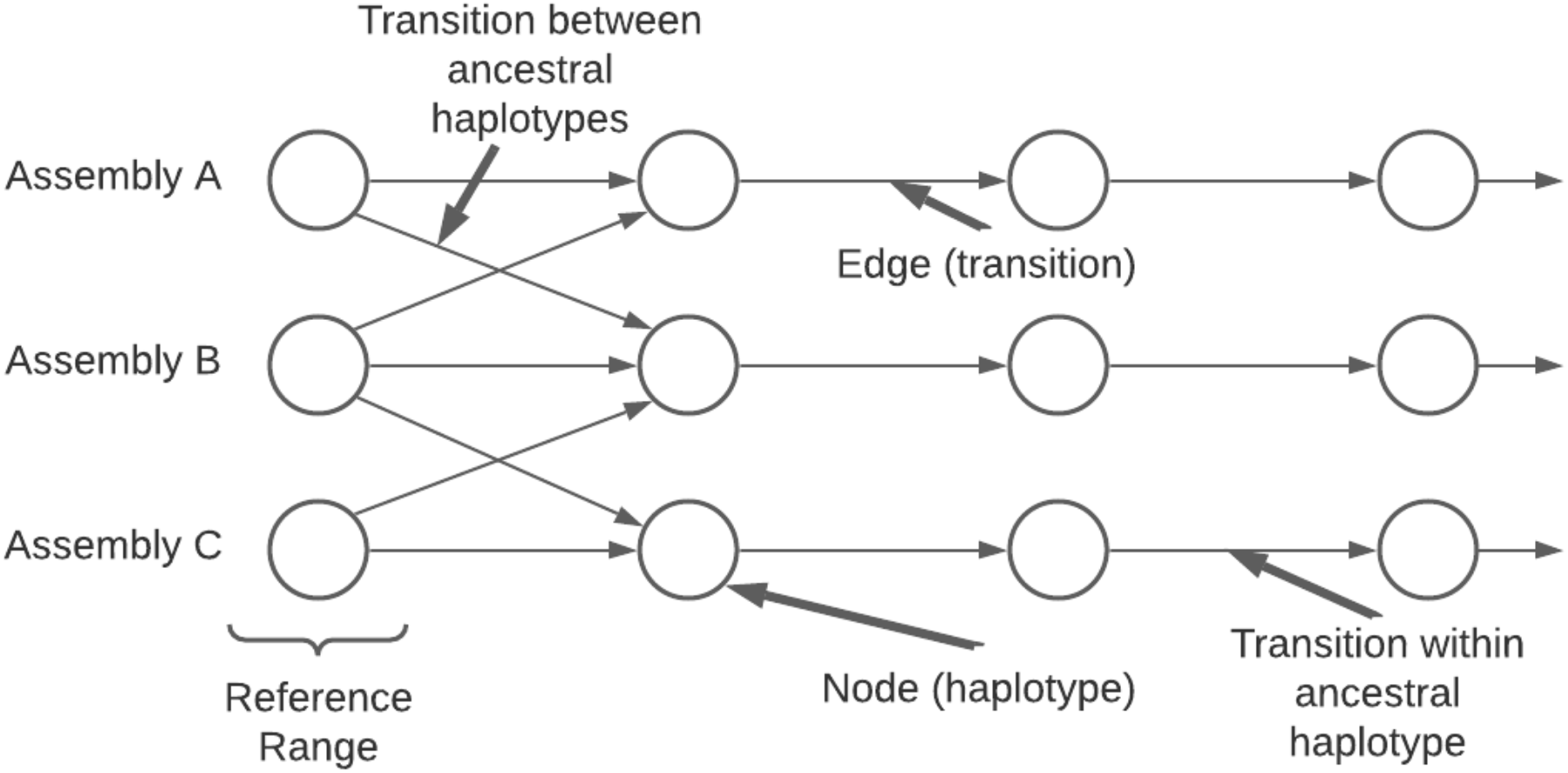
Diagram of a Practical Haplotype Graph (PHG). Imputation uses emission probabilities from the nodes and transition probabilities from the edges.

The PHG database (Supplementary Figure 1) stores all the data necessary to build a haplotype graph, which exists only in computer memory. The data behind the graph is persistent, while the graph is not. The first step in creating a PHG database defines the reference range endpoints and assigns reference ranges to user-defined groups. One useful strategy defines the endpoints from gene boundaries. The reference ranges containing genes are assigned to one group and the intergenic ranges to another group, which together cover the entire genome. After that, the reference genome is loaded followed by haplotypes from other individuals.

The basic steps to populate the PHG database include importing the reference range endpoints, loading the reference sequence, creating additional haplotypes, and making consensus haplotypes. Figure 2 provides an overview of the process used to populate the database after the reference genome and reference ranges have been set up. The additional haplotypes can be created from assemblies by aligning those to the reference genome or from whole genome sequence (WGS). Haplotypes should be loaded from enough individuals to populate the database with the common haplotypes present in the target population. Once the haplotypes have been loaded, skim sequence can be used to impute additional genotypes. The resulting information is stored in the database in a compressed format.

**Figure 2.**
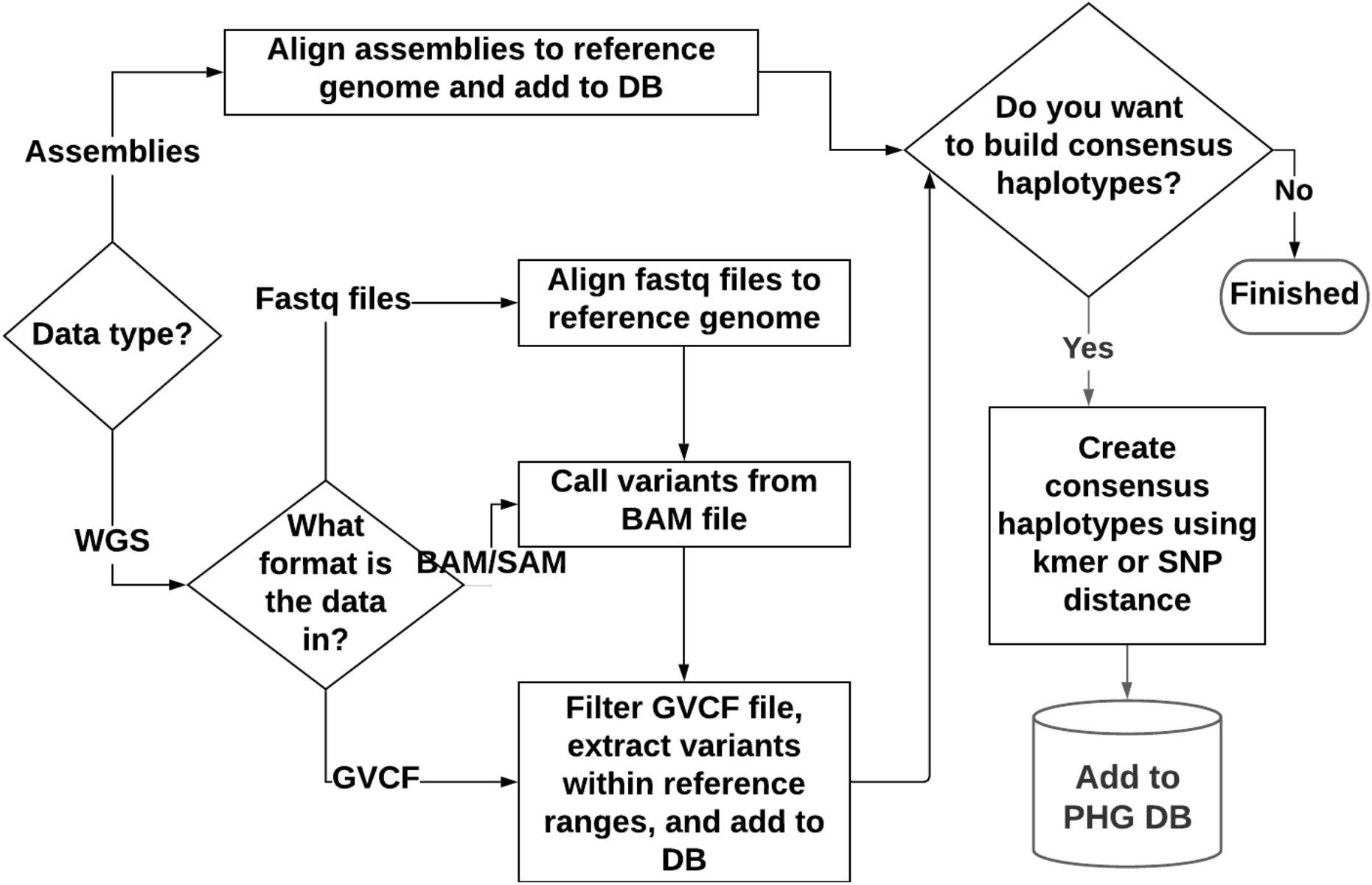
Building a PHG database

The process of creating haplotypes involves aligning sequence to the reference genome, but the method and results are different depending on whether assembled genomes or WGS are used. In the case of WGS, only sequence that aligns well to a single location on the reference genome is used to create haplotypes. For many species, such as maize, WGS data provides good haplotype coverage for genic reference ranges but not for many of the intergenic ranges, which often do not align well. For example, in any two maize inbreds, large segments of intergenic ranges can carry completely different sequences. With an assembly, if two adjacent genes can be aligned to reference, the intervening sequence can be assigned to the intergenic reference range even if the intergenic sequence itself does not align, but with WGS unaligned sequence is not assigned to a haplotype.

Once a PHG database has been populated with haplotypes, it can be used to impute genotypes from skim sequences or SNPs (Supplementary Figure 2). Any new sample can be represented as a collection of haplotypes already present in the PHG. The model used for imputation is very similar to the Li and Stephens model (Li and Stephens, 2003), which views genotypes as composed of segments from reference or ancestral haplotypes.

The method used by the PHG for imputation differs from most other imputation software in one key way. It can use either SNP calls or raw sequence reads as input. Most other software starts with SNPs, often considered to be assayed with low error. PHG imputation can start with raw sequence reads and use those directly to tag haplotypes. It does so by aligning those reads to the PHG pangenome, identifying which haplotypes match each read, then using that information together with the information about which nearby haplotypes appear on the same chromosome to determine which set of haplotypes is most likely to have generated the observed reads. Alternatively, SNPs from any genotyping method can be used to tag haplotypes. A hidden Markov model (HMM) uses the resulting read mapping counts to impute the most likely path through the haplotype graph. The results are stored as a list of haplotype ids. More information about the alignment software used to build the PHG and to impute paths, imputation details, and the database schema can be found in the supplementary methods.

### Software Architecture and Distribution

The PHG software is a combination of custom code written using Java and Kotlin (a modern JVM language), third-party software, and a set of helper scripts. The main PHG functions have been organized as plugins based on the TASSEL 5 plugin architecture as described on the TASSEL wiki (https://bitbucket.org/tasseladmin/tassel-5-source/wiki/Home). The primary reasons for using the plugin architecture are that it provides a consistent API, which can be called from a command line or from the TASSEL GUI, and is self-documenting. The custom code is freely available, open source, and distributed with TASSEL 5 as phg.jar.

All of the steps in the PHG pipeline can be run with individual plugins from a command line. Because running the PHG with a production database typically requires a large amount of RAM (> 50GB for some steps) and uses multiple CPUs, it has been designed to run on a server using terminal commands. The database connection parameters must be stored in a configuration file. Parameters for individual plugins can be specified on the command line or included in the configuration file. Because the pipeline is complex and overall has more than one hundred user-settable parameters, it has been organized into a few pipeline plugins covering the entire process from building the database, populating it with haplotypes, and imputing genotypes. The pipeline plugins, individual components and sample configuration files are all documented at the PHG wiki (https://bitbucket.org/bucklerlab/practicalhaplotypegraph/wiki/Home) along with directions for creating an example database for testing (https://bitbucket.org/bucklerlab/practicalhaplotypegraph/wiki/UserInstructions/ExampleDatabase.md).

The PHG is deployed as a Docker image. Each Docker image contains a snapshot at a particular point in time of the software used for processing. As new versions of a dockerized image are created, the older versions remain available on docker hub (https://hub.docker.com/r/maizegenetics/phg). This makes it easy for users to replicate analyses using a specific version of the complete software environment.

In addition to the command line API, we provide two additional API’s. An alternative interface to the PHG that provides useful methods for examining and analyzing the contents of a PHG database is rPHG, implemented as a package for the statistical programming language R. PHG objects are built and stored in the R environment using the PHG API along with a Java to R interface (https://CRAN.R-project.org/package=rJava). Details about downloading and running rPHG as well as source code can be found on Bitbucket (https://bitbucket.org/bucklerlab/rphg/wiki/Home).

Another API available with the PHG is a Breeding API (BrAPI) (Selby *et al*., 2019) compliant web service written using Ktor (https://ktor.io). Ktor is a framework for creating web applications using the Kotlin programming language. BrAPI is a specification for a RESTful web service for sharing plant breeding data. Specifically, the PHG web server implements the subset of BrAPI calls that support genotype information. Any breeding or genetics application that implements the BrAPI V2 standard for genotypes will be able to access data from PHG databases connected to a web service. Because the PHG BrAPI web server is under active development, no public server instances are available at this time.

## Results and Discussion

Important features of a pangenome representation include efficient data compression and the accuracy of derived analyses like imputation. Using simulated data in maize, we evaluate these two characteristics of the platform. Empirical studies are already available using earlier versions of the pipeline (Jensen *et al*., 2020; Valdes Franco *et al*., 2020; Jordan *et al*., 2021; Long *et al*., 2021). To evaluate performance, we use the maize PHG with 26 high quality assembly genomes and low density GBS data on 5,000 recombinant inbred lines derived from those lines. The maize reference genome is 2.3Gbp in size.

### Alignment Performance

The initial building of the PHG from assemblies requires the alignment of high-quality genomes to the reference genome. Adding new assemblies to the PHG takes 7-8 CPU hours with two-thirds of the time used by the MUMmer4 alignment process. While this whole process can be parallelized, it is limited by the speed and sensitivity of tools like MUMmer.

The PHG still relies on standard high-performance aligners such as minimap2 in order to map reads for path imputation. To estimate the speed and memory requirements, we compared reference genome (2.3Gbp reference) alignment with alignment to a PHG with 26 genomes. The RAM requirements increased by a factor of 4.4 from 34Gb to 150Gb, while the processing time only increased 2.3-fold. While the memory requirements are substantial, processing is efficient given the 26-fold increase in the search space.

### Data Compression

A key aim of the PHG database is to efficiently store and represent genomes in terms of disk and in-memory space. Most of the storage space used by the PHG database consists of four types of data: haplotype sequence, haplotype variants, read mapping counts, and imputed paths. The data types vary greatly in storage space use as compared to alternative formats for similar data.

#### Haplotype sequence

Haplotype sequence for assemblies is stored as compressed strings; the amount of disk space used is similar to compressed FASTA files for the same assemblies.

#### Haplotype variants

The variant data for maize assemblies aligned with MUMmer4 uses about 80% as much space as the files output by MUMmer4.

#### Read mapping counts

The space used by read mappings varies depending on the type and coverage of the data used to generate them as well as the number of haplotypes to which they were aligned. Aligning 30 million simulated reads (3Gb Fastq file) against a 26-genome maize PHG would normally produce a 50Gb BAM file. However, recording only the counts of best haplotypes combinations results in a 69Mb file (43-fold smaller than initial fastq, and 724-fold smaller than the BAM file).

#### Imputed path

A single imputed path uses about 250KB of database storage, whereas a gzipped gvcf generated from homozygous PHG paths for a maize PHG requires about 176Mb per sample, a 704-fold difference. A major advantage of the PHG is that it can store a large number of fully imputed genomes very efficiently. In large panels of germplasm such as 4,705 NAM RILs, even if we only write the SNPs to a standard VCF file, the haplotype representation is roughly 1000-fold smaller.

### Haplotype Mapping Accuracy

Imputing haplotypes and SNPs for a large number of samples using a pangenome containing related germplasm is a central goal of the PHG. PHG imputation performance requires that sequencing reads are accurately mapped to haplotypes in the PHG. While short read alignment accuracy is a problem for all genomics, it can be a larger problem in massive pangenomes with redundant low copy sequences. To evaluate mapping performance, we simulated reads generated from haplotype sequence stored in the PHG database for the inbred NAM parents B73, CML247, and Oh43. After those reads were mapped back to all 26 genomes in the database, we asked whether each read mapped to the sequence from which it came.

Because most reads align to more than one haplotype, if the read mapped to the haplotype of the line from which it came, it was counted as a correct mapping even if it mapped to other haplotypes as well. A read that did not map to the haplotype from which it was drawn counted as mapping incorrectly. Testing whether reads generated from one of the assemblies map back to the originating assembly is an important test of a pangenome system.

We found accuracy was most sensitive to read types, how haplotypes were collapsed, and minimizer redundancy. Specifically, using paired-end reads improved accuracy three to sixfold over single-end reads across a range of collapsing approaches (Figure 3). This increase in accuracy is the result of paired-end reads mapping more accurately to paralogous retrotransposons, as the advantage of paired-end reads was smaller in genic regions (Supplementary Figure 3) than in the genome as a whole. Increasing minimizer redundancy improved the accuracy of single-end reads more than the accuracy of pair-end reads, again because of improvement in mapping to repetitive regions (Figure 3).

**Figure 3.**
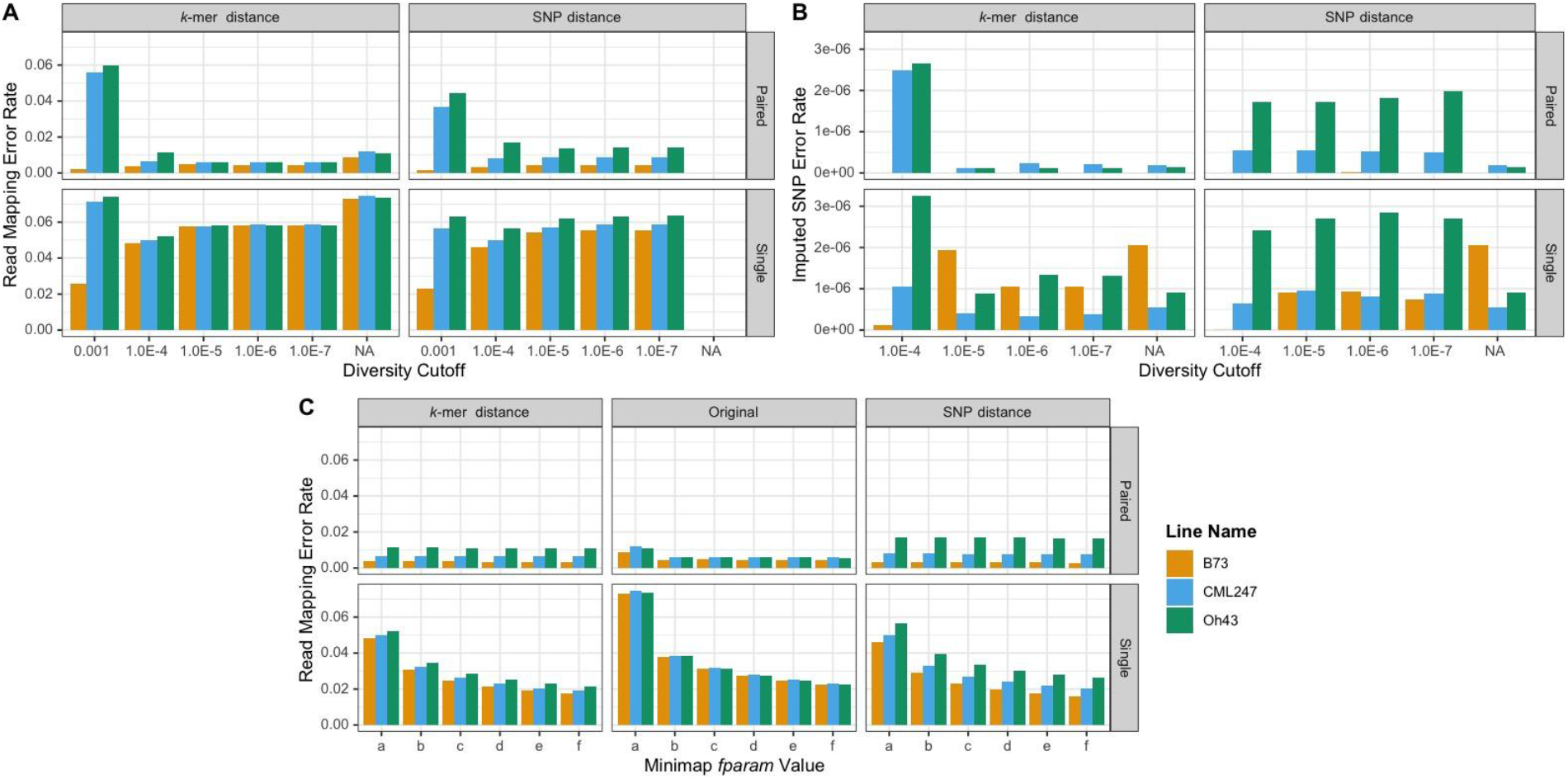
Effect of parameters on accuracy. A) Read mapping error rate for the whole genome as a function of the maximum diversity (mxDiv) parameter for determining consensus haplotypes, read type (paired-end, single), and distance matrix method (kmer, SNP). Read mapping error rate is the number of reads not mapping to the target haplotype divided by the total number of reads. NA labels the result of mapping against the original, non-consensus haplotypes. B) Imputed SNP error rate for paired and single-end reads for different values of the diversity cutoff and consensus method. Error rate equals the number of wrong SNP calls divided by the number of base pairs of sequence. Where the B73 bar is absent, the error rate was zero. C) Read mapping error as a function of minimizer redundancy controlled by the minimap2 f-parameter. f parameter values are a) f1000,5000 [default]; b) f5000,6000; c) f10000,11000; d)15000,16000; e) f20000,21000; f) f25000,26000. mxDiv = 1e-4.

The size of the entire pangenome can be reduced by collapsing similar haplotypes. Haplotypes can be collapsed using divergence measured either by SNP or kmer similarity. The kmer based approach, which is alignment free but slower, detects both SNP and indel differences between haplotypes. Overall the kmer based approach was more robust and decreased mapping error by roughly 1.7% across the entire genome (Figure 3) and 15.8% in genic regions (Supplementary Figure 3). While more extreme collapsing of haplotypes (mxDiv=0.001) increased error, the pipeline was insensitive to changes when mxDiv was below 0.00001. There was substantial bias at high levels of collapsing (mxDiv=0.001) because of the way a haplotype is chosen to represent a cluster. The assemblies in the database are ranked by quality and the best quality haplotype is chosen to represent a cluster. For the simulation, the rank order was B73 > CML247 > Oh43, which resulted in better accuracy for the higher ranked assembly (B73). With SNP based distance, but not kmer based distance, that bias persisted at lower collapsing levels because haplotypes with no SNPs but with indels were collapsed whereas with kmer similarity only identical haplotypes were collapsed. As a result, kmer based distance should be the preferred method for collapsing haplotypes.

Alignment accuracy was also impacted by tuning the way the aligner (minimap2) handles highly repetitive sequences. Aligners frequently purge highly repetitive kmers or minimizers from their lookup tables in order to maintain performance. This can be problematic in pangenomes. For the data presented here, we aligned against 26 genomes at the same time. Not only was the sequence repeated within genomes, it was repeated across them as well. In minimap2, the f-parameter determines which minimizers are used for alignment. When f has an integer value, then minimizers with more than f occurrences are not used. Because the default f-parameter was not optimal, mapping accuracy could be improved by nearly twofold with higher f-parameters settings (Figure 3). For paired-end reads with collapsed haplotypes, the default f-parameter was sufficient. Overall, read mapping errors were minimized with kmer based collapsing, an increase in the f-parameter, and use of paired end sequencing.

### Haplotype Imputation Accuracy

For a set of read mapping counts, a hidden Markov model (HMM) finds the path through a haplotype graph that best explains the observed data. We tested the impact of the read type, minimap’s f-parameter, and haplotype collapsing approach on the haplotype error rate (Supplementary Figures 4 & 5). Haplotype error rates were about sevenfold lower on average than the mapping error rates, which shows that the HMM model corrects many of the mismapping errors. Paired-end reads performed better than single end reads, but changes in f-parameter, diversity cutoff, and distance method had little impact. The increased error rate for non-consensus haplotypes compared to consensus haplotypes is misleading because, prior to collapsing, the HMM may have chosen a haplotype that was identical to the read source but from a different parent taxon. Overall, the HMM identified haplotypes with error rates as low as 0.1%.

While reconstructing the ancestral haplotype is useful in closed breeding pools and for some applications, generally users are most interested in imputing variants correctly. At the SNP level, the error rates approach one in a million under a range of realistic conditions for read types and haplotype collapsing (Figure 3). In a closed population as simulated here, error rates approach zero if the PHG is tuned appropriately and using appropriate read types. While validating the platform and providing insight into the parameters, this simulation did not test the effect of genotyping errors or imputing individuals not used to build the database. These issues were addressed by evaluating the PHG with real data.

### Empirical Studies

Other studies describe and evaluate the use of a PHG for sorghum (Jensen et al., 2020), maize (Valdes Franco et al., 2020), wheat (Jordan et al., 2021), and cassava (Long et al., 2021). The sorghum project used WGS data for 24 inbred lines at an average 8x coverage to populate a database with haplotypes. Imputing a set of lines mostly derived from the 24 inbred lines produced SNP calls that were nearly the same as a GBS-specific pipeline and as effective for genomic selection. The maize study used the same database that was used for the simulation reported here. Genotypes were imputed from GBS sequence reads for over 5,000 NAM recombinant inbred lines (RILs) resulting in an average genotype error rate less than 0.01. The wheat PHG was built from the exome capture data of 65 diverse wheat lines. Exome capture, GBS, and skim sequence data was used to accurately impute the genotypes of a panel of lines. The cassava paper used haplotypes derived from runs of homozygosity in otherwise heterozygous cassava lines to populate a PHG and showed that the resulting PHG could be successfully used to impute genotypes from 1X coverage skim sequence. Taken together these studies show that the PHG can be used with a variety of data sources and species.

## Conclusions and Future Directions

The PHG is currently a powerful tool for storage, retrieval, and imputation of haplotypes that can be used for both genomics and breeding. It is already being used to merge the results of diverse genotyping platforms to construct unified databases for entire species. Specifically, the PHG will be a useful tool for breeding programs for merging historic genotyping data generated from multiple methods, for switching genotyping to low cost providers, and for tracking haplotype selection across generations. A current limitation is that accuracy of particular portions of paths is not estimated, which is especially important when samples with new haplotypes are being analyzed. Statistics derived from the alignment and HMMs in conjunction with machine learning should be able to estimate path accuracy for individual reference ranges.

The “practical” in PHG refers to the decision not to construct a complete variant graph because that construction is very difficult for genomes with large numbers of active retrotransposons. Instead, in the case of assemblies the retrotransposons are assigned reference positions based on nearby alignable sequence or, for WGS-based haplotypes, simply not used when they cannot be aligned. Additionally, since recombination is substantially focused within genes or near their open chromatin flanks, a haplotype graph divided at the edges of shared genes provides a reasonable compromise between capturing ancestral haplotypes and recombination. Finally, this practical representation allows the use of powerful aligners such as minimap2 to impute PHG paths from short or long reads. In the future, we see the potential to evolve the PHG into a graph of graphs by modeling each reference range with a variant graph (Garrison *et al*., 2018).

In the plant breeding community, BrAPI (Selby *et al*., 2019) is becoming widely accepted as a standard for sharing genotypic and phenotypic data across systems and analysis tools. Because the BrAPI standard for sharing large genomic dataset is under development and because the PHG DB can serve as a repository for genomic data with up to 1000-fold data compression, we are working with the community to improve variant and haplotype graph support in BrAPI version 3.

Visualization and tools to work with haplotype graphs are still in their infancy, but as APIs become more robust and standardized, a wide range of tools in bioinformatic languages such as R and Python will need to be developed. The development of tools that allow diversity to be accurately represented and functional variation to be identified can help drive both basic and applied research into natural variation.

## Acknowledgements

This material is based upon work supported by the USDA-ARS, NSF Research-PGR Grant No. IOS-1238014 and IOS-1822330, and the Bill and Melinda Gates Foundation (OPP1159867 and OPP1175661).

## References

Baucom, R.S. et al. (2009) Exceptional diversity, non-random distribution, and rapid evolution of retroelements in the B73 maize genome. PLoS Genet., 5, e1000732.

Bennetzen, J.L. (2000) Transposable element contributions to plant gene and genome evolution. Plant Mol. Biol., 42, 251–269.

Brunner, S. et al. (2005) Evolution of DNA sequence nonhomologies among maize inbreds. Plant Cell, 17, 343–360.

Eizenga, J.M. et al. (2020) Pangenome Graphs. Annu. Rev. Genomics Hum. Genet., 21, 139–162.

Garrison, E. et al. (2018) Variation graph toolkit improves read mapping by representing genetic variation in the reference. Nat. Biotechnol., 36, 875–879.

Jensen, S.E. et al. (2020) A sorghum practical haplotype graph facilitates genome-wide imputation and cost-effective genomic prediction. Plant Genome, 13, e20009.

Jordan, K.W. et al. (2021) Development of the Wheat Practical Haplotype Graph Database as a Resource for Genotyping Data Storage and Genotype Imputation. bioRxiv, 2021.06.10.447944.

Li, N. and Stephens, M. (2003) Modeling linkage disequilibrium and identifying recombination hotspots using single-nucleotide polymorphism data. Genetics, 165, 2213–2233.

Li, R. et al. (2010) Building the sequence map of the human pan-genome. Nat. Biotechnol., 28, 57–63.

Llamas, B. et al. (2019) A strategy for building and using a human reference pangenome. F1000Research, 8, 1751.

Long, E.M. et al. (2021) Genome-wide Imputation Using the Practical Haplotype Graph in the Heterozygous Crop Cassava. bioRxiv, 2021.05.12.443913.

Rakocevic, G. et al. (2019) Fast and accurate genomic analyses using genome graphs. Nat. Genet., 51, 354–362.

Rodgers-Melnick, E. et al. (2015) Recombination in diverse maize is stable, predictable, and associated with genetic load. Proc. Natl. Acad. Sci. U. S. A., 112, 3823–3828.

Schnable, P.S. et al. (1998) Genetic recombination in plants. Curr. Opin. Plant Biol., 1, 123–129.

Selby, P. et al. (2019) BrAPI—an application programming interface for plant breeding applications. Bioinformatics, 35, 4147–4155.

Sherman, R.M. et al. (2019) Assembly of a pan-genome from deep sequencing of 910 humans of African descent. Nat. Genet., 51, 30–35.

Sun, S. et al. (2018) Extensive intraspecific gene order and gene structural variations between Mo17 and other maize genomes. Nature Genetics, 50, 1289–1295.

Valdes Franco, J.A. et al. (2020) A Maize Practical Haplotype Graph Leverages Diverse NAM Assemblies. bioRxiv, 2020.08.31.268425.

