## Supplemental Methods and Figures for "The Practical Haplotype Graph, a platform for storing and using pangenomes for imputation"

### PHG Supplementary Material

#### Supplementary Figures

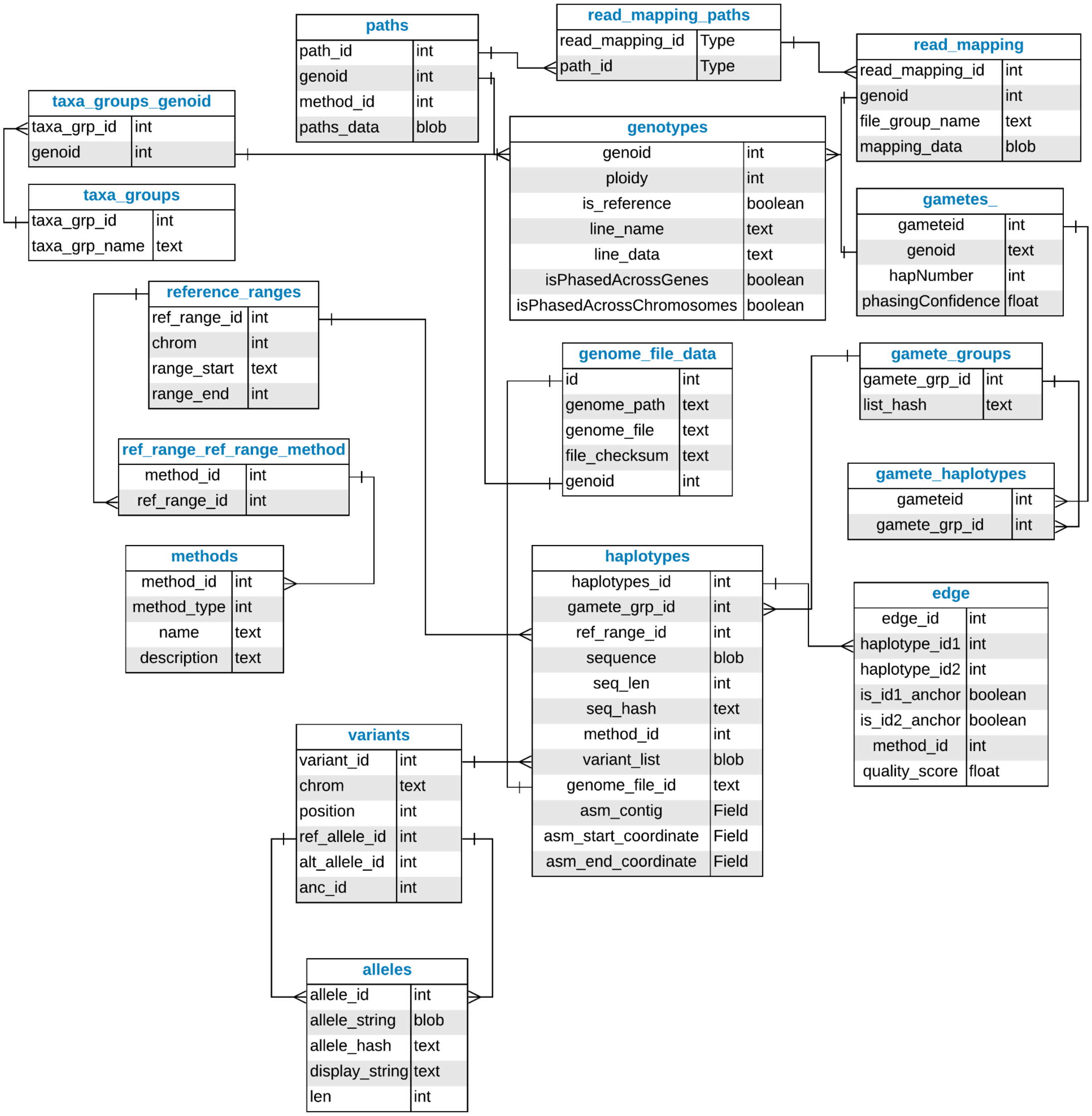


Supplementary Figure 1. PHG database schema

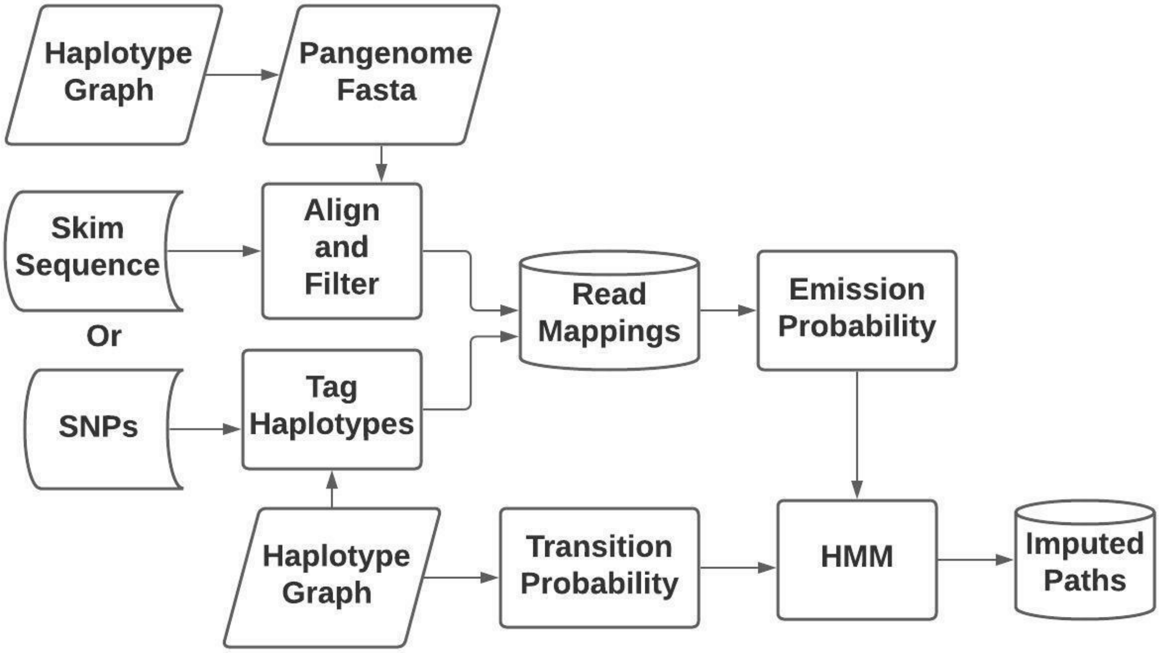


Supplementary Figure 2. Flow chart for PHG imputation. Skim sequence or SNPs tag haplotypes to produce read mappings that are stored in the PHG DB. The read mappings and a haplotype graph provide the probabilities to impute paths through the graph using a hidden Markov model (HMM)

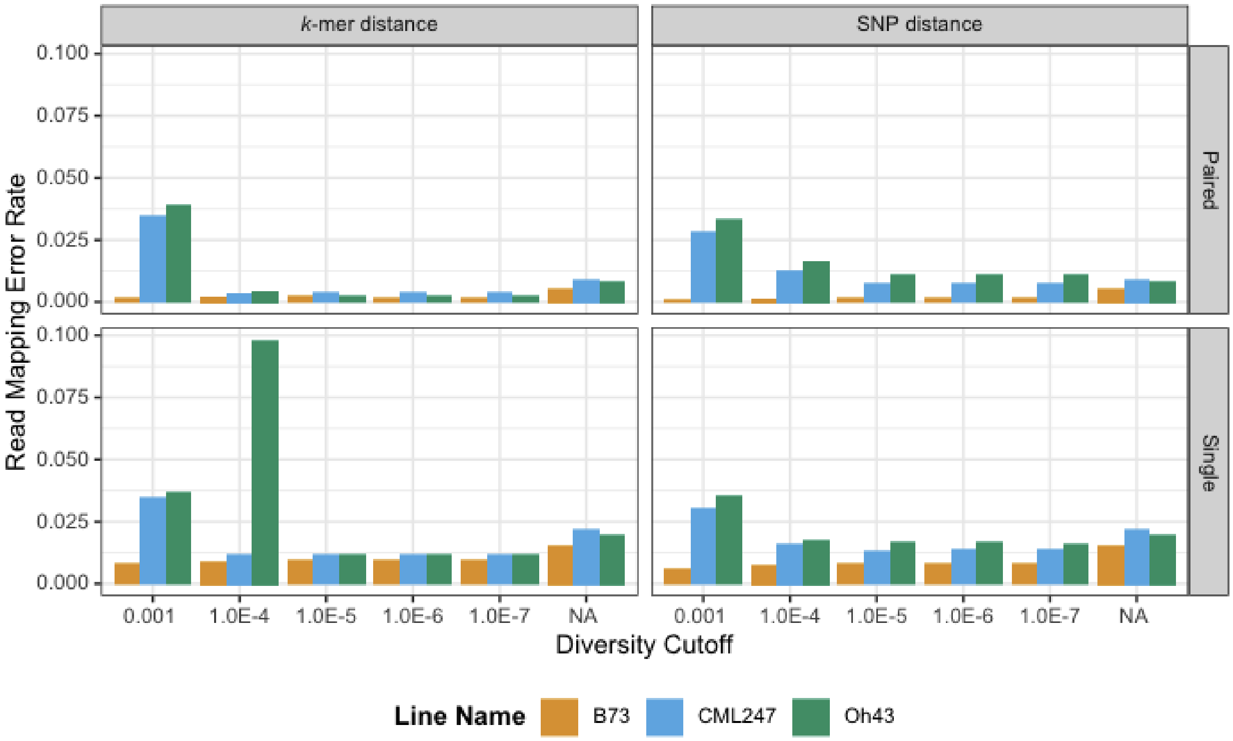


Supplementary Figure 3. Read mapping error rate for genic reference ranges as a function of the diversity cutoff (mxDiv) parameter for determining consensus haplotypes, read type (paired-end, single), and distance method (kmer, SNP). NA labels the result of mapping the original, non-consensus reads.

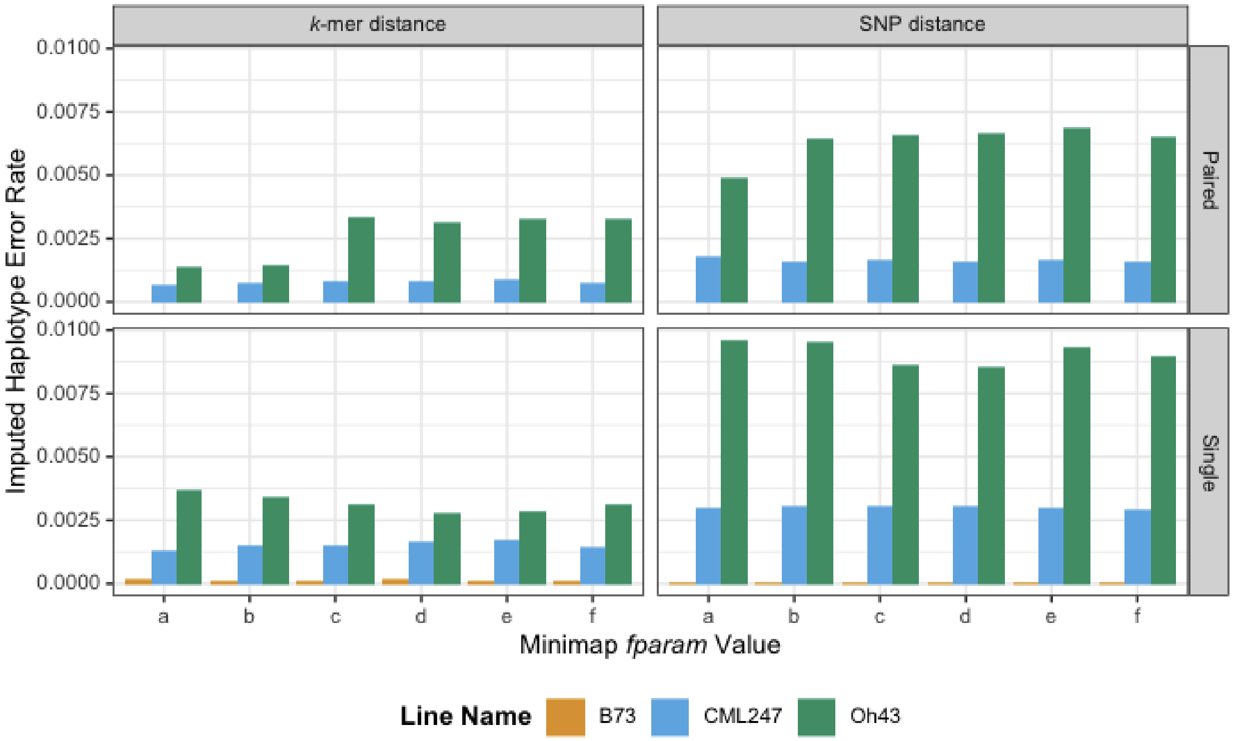


Supplementary Figure 4. Haplotype error rate as a function of the Minimap2 f parameter (fParam parameter), read type (paired-end, single), and distance matrix method (kmer, SNP), where a = f1000,5000, b = f5000,6000, c = f10000,11000, d= f15000,16000, e = f20000,21000, and f = f25000,26000. The analysis used mxDiv = 1e-4. Where there is no red bar, the error rate for B73 was 0.

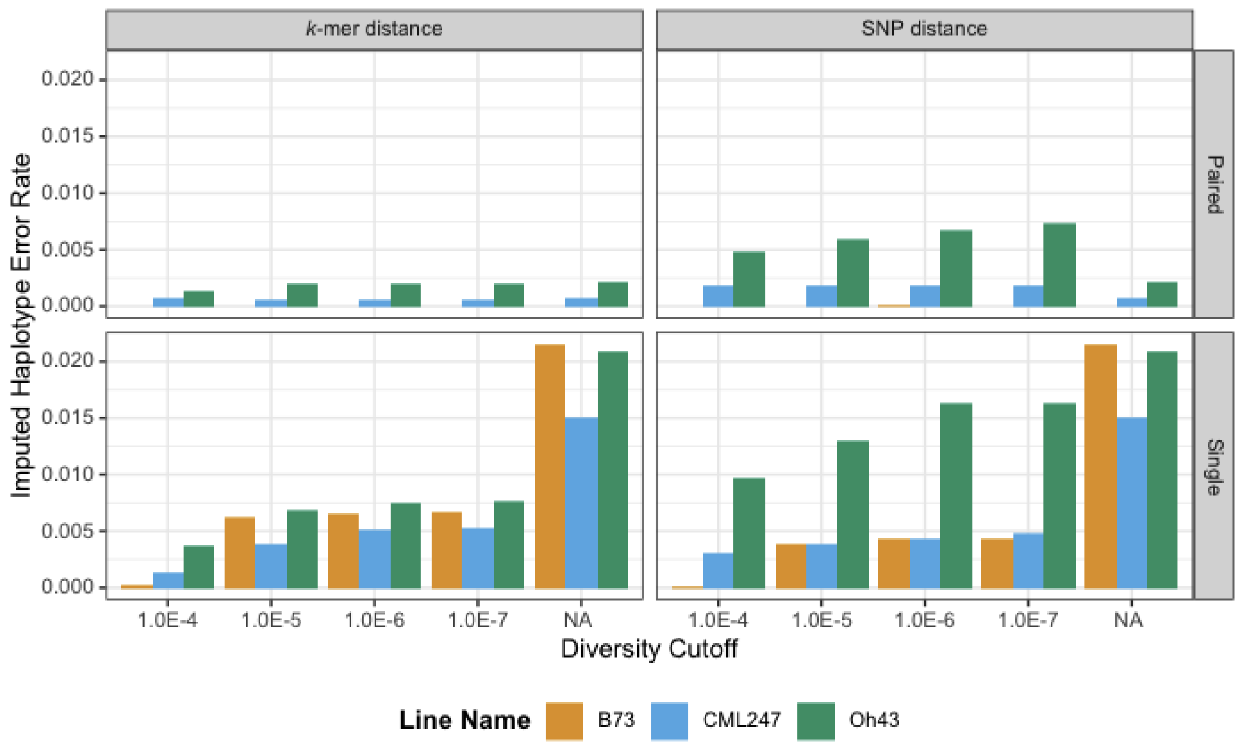


Supplementary Figure 5. Haplotype error rate as function of maximum diversity (mxDiv parameter), read type (paired-end, single), and distance matrix method (kmer, SNP). Diversity cutoff = NA denotes non-consensus haplotypes.

### Supplementary Methods

#### Alignment software

The PHG uses different alignment software for different tasks in the pipeline. Assemblies are aligned to the reference genome using Mummer4 (Marçais et al., 2018) then a longest path algorithm assigns specific genome segments to reference ranges. When WGS is used to add haplotypes to the database, it is first aligned to the reference genome using bwa mem (Li, 2013). The output of that step is used by GATK HaplotypeCaller (Van der Auwera and O'Connor, 2020) to produce a GVCF file that is loaded to the database. Users can choose to substitute their own pipeline to create BAM files or GVCF files instead. The first part of the imputation pipeline uses minimap2 (Li, 2018) to align sequence to the PHG pangenome to generate read mapping counts.

#### Imputation Details

Imputation using the PHG has two distinct steps. The first is generating read mapping counts, which are stored in the database. The second is using those counts together with a haplotype graph and an HMM to impute the most likely path through the graph. Read mapping counts can be created from sequence or from SNP calls. To generate read mappings, the sequence from a haplotype graph is written to a fasta file. Each haplotype in the graph produces a record in the fasta file. As a result, when minimap2 aligns a read, it returns information for each matching haplotype. The haplotypes with the lowest edit distance for a single read form a haplotype set. The read mapping counts stored for a single sample is a list of all the haplotype sets for that sample together with a count of times they occurred. Read mapping counts can also be generated from a VCF file of SNP calls.

The second step uses read mapping counts to derive the most likely path through the graph. A path through the haplotype graph can be represented as a list of haplotypes and defines an imputed genome. The read mapping counts are used to calculate emission probabilities while the haplotype graph is used to calculate transition probabilities. These probabilities are the input required by the HMM. The algorithms available to solve the HMM are the Viterbi algorithm and the forward-backward algorithm (Rabiner, 1989). The Viterbi algorithm finds the most likely path given the input data. The forward-backward algorithm calculates the likelihood for each individual haplotype and determines a path by choosing the most likely node in each reference range. The PHG has separate code for imputing homozygous (effectively haploid) and heterozygous diploid individuals (Supplementary Figure 2).

Database

The PHG uses a relational database, either PostgreSQL or SQLite, to store data for the pangenome graph. An SQLite database consists of a single file, requires minimal setup, and can be copied and moved easily, which makes it a good platform for development and testing. PostgreSQL is an industry standard with high scalability, making it suitable for large databases in a production environment. Both are open source options and provide for user flexibility. If using PostgreSQL, the database may be local or remote.

Occasionally, the database schema changes and new versions of the PHG software may be incompatible with older versions of the database. To avoid forcing users to rebuild databases, which can be a time consuming process, Liquibase (https://www.liquibase.org/) is used to update older databases. The PHG pipeline plugins automatically compare the database and software versions and update the database when needed. Alternatively, users can download and run the phg_liquibase docker to perform an update manually.

The PHG database schema (Supplementary Figure 1) describes how data is stored. Genotypes, gametes, and gamete groups are used to store and manage taxon (sample) names. Gamete groups allow a single consensus haplotype to be assigned to multiple genotypes. Gametes provide a way to store more than one haplotype per genotype.

The haplotypes, variants, and alleles tables store information about haplotypes. Because the same allele or variant may appear multiple times for different haplotypes, alleles and variants have their own tables. This reduces data redundancy and improves memory efficiency. Haplotype sequence is stored as compressed strings. Variant lists are stored as an encoded and compressed list of variant ids. Doing so reduces storage space but requires data to be processed after retrieving it from the database in order to be used.

Simulated Data

The results described in the discussion section were obtained by analyzing reads simulated from haplotype sequence stored in a maize database populated with the maize NAM parent assemblies (Valdes Franco et al., 2020). Single reads were created by selecting 150 base pair segments from each haplotype at random start sites for inbred lines B73, CML247, and Oh43. Paired-end reads were created by selecting two 150 base pair segments separated by 50 base pairs (making a 350 bp fragment) at random start sites and reverse-complementing the second read of the pair. The total length of sequence selected for each reference range and read type equaled 0.1 of the haplotype length to approximate 0.1X coverage. The resulting simulated reads were written to FASTQ files to use as input for imputation. The script that ran the analysis of the simulation was written in Kotlin, while the code that generated the graphs was written in R as an R notebook. Both are stored in the public bitbucket PHG repository. The analysis itself was run using the docker maizegenetics/phg:0.0.27.

Alignment speed estimation:

To test the memory footprint and speed of aligning short reads to a PHG pangenome, we created a FASTQ file containing 30 million GBS reads taken from NAM recombinant inbred lines. Each read was 150 base pairs. We extracted the haplotype sequences of the NAM parent assemblies from the database used for simulation and indexed the resulting fasta using minimap2's -sr presets. We also indexed the maize reference genome (B73 AGPv5) using the same parameters. Note that the maize reference genome is one of the NAM founder assemblies. We then aligned our test GBS FASTQ against both of our indices keeping track of time and RAM usage. We set the number of threads parameter (-t) to 20 and number of matches parameter (-N) to 50 for both alignments. We also piped the output of minimap2 into a BAM file using samtools.


Li,H. (2013) Aligning sequence reads, clone sequences and assembly contigs with BWA-MEM. *arXiv [q-bio.GN]*.

Li,H. (2018) Minimap2: pairwise alignment for nucleotide sequences. *Bioinformatics*, **34**, 3094–3100.

Marçais,G. *et al.* (2018) MUMmer4: A fast and versatile genome alignment system. *PLoS Comput. Biol.*, **14**, e1005944.

Rabiner,L.R. (1989) A tutorial on hidden Markov models and selected applications in speech recognition. *Proc. IEEE*, **77**, 257–286.

Valdes Franco,J.A. *et al.* (2020) A Maize Practical Haplotype Graph Leverages Diverse NAM Assemblies. *bioRxiv*, 2020.08.31.268425.

Van der Auwera,G.A. and O’Connor,B.D. (2020) Genomics in the Cloud: Using Docker, GATK, and WDL in Terra ‘O’Reilly Media, Inc.’
